# An innovative tool for non-invasive contact-free pathogen monitoring in animal saliva

**DOI:** 10.64898/2026.02.11.705368

**Authors:** Maristela Martins de Camargo, Aspen Caim Fernandes Gonçalves, Diana Netto Hernandez Blazquez, Maggie E. Weber, Washington Agostinho, Zubin Kremer Guha, Melissa Sá Fernandes, Mariana Furtado, Aaron Dollar, Paulo Brandão, Serap Aksoy, Vanessa O. Ezenwa, Dawn Zimmerman

## Abstract

Habitat fragmentation, climate change, poaching, human-wildlife conflicts, and infectious diseases are the main threats to biodiversity conservation. They alter host-pathogen dynamics, reduce viable conservation areas, and promote genetic isolation, resulting in physiological stress among animal populations. Moreover, increased proximity between domestic and wild animals further facilitates disease spillovers exposing naïve host species and ecosystems to new pathogens. Of the more than 200 known zoonotic diseases, approximately 60% originate from animals, contributing significantly to the global infectious disease burden. Here, we describe the development of an innovative non-invasive approach for biological sampling that has been validated in mice and shelter cats. Our device consists of a disposable plastic cassette that through odor attractants lures animals to lick a filter paper. This saliva collection approach allowed for the detection of RNA viruses by RTqPCR and third-generation sequencing. RTqPCR oral swab and licked paper results showed that both methods significantly predicted serological status. Our sequencing results revealed the richness of the gene space, demonstrating the potential of this device for discovering rare or unknown species circulating in the saliva donor, enabling this player to be recognized as an environmental sentinel. This study demonstrates the feasibility of deploying this device in sheltered/captive animal settings as well as under laboratory simulations of different environments, providing necessary foundations for future field applications. Our methodology holds great potential for monitoring zoonotic pathogens in both captive and free-ranging animals, to even possibly allow proactive mitigation measures prior to spillover, without interfering with the natural animal behaviour and social structures.

**Visual abstract:** 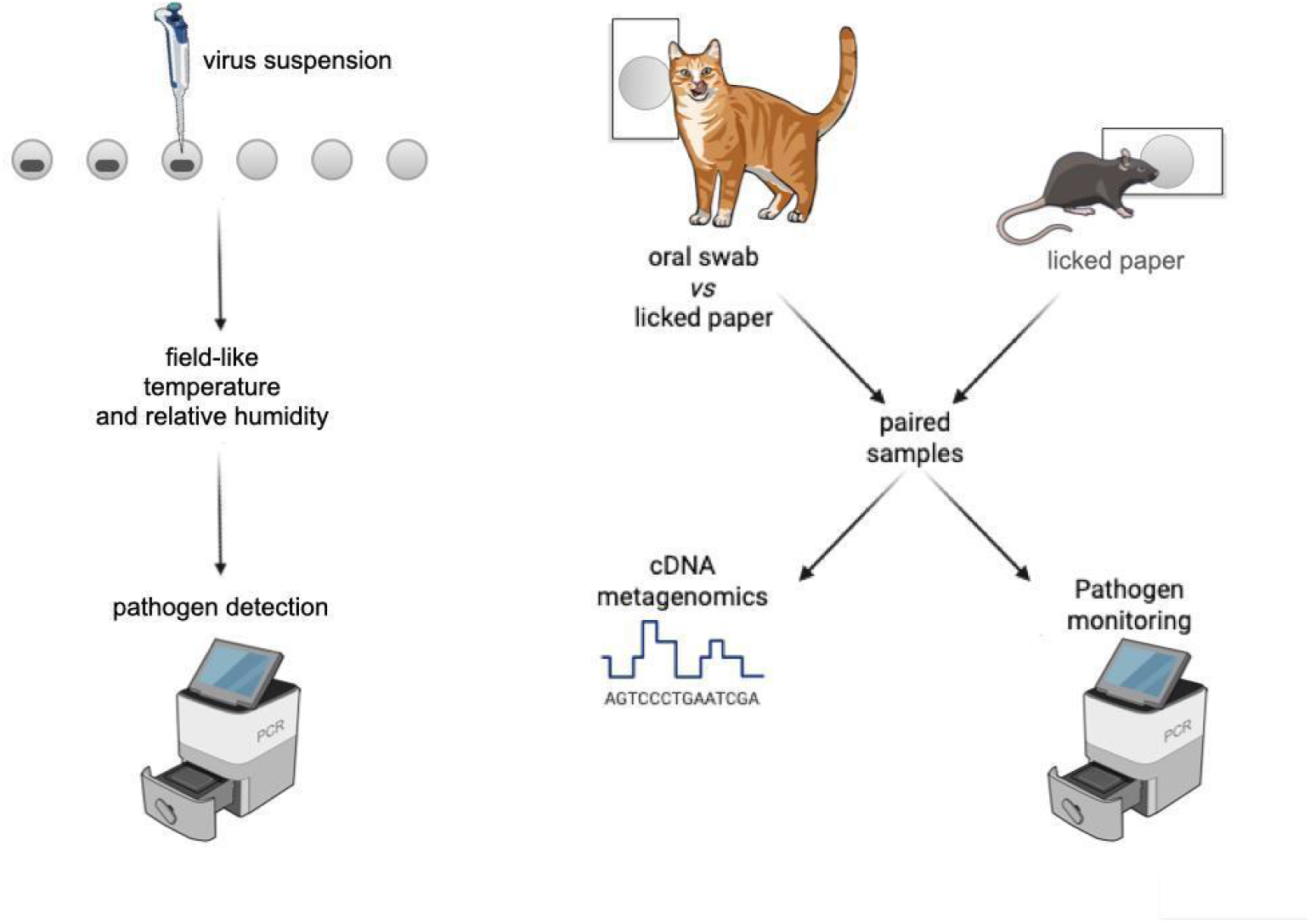

## Introduction

Habitat fragmentation, climate change, poaching, human-wildlife conflicts, and infectious diseases are among the main threats to biodiversity conservation.^1,2^ These threats, coupled with an increasing human-wildlife interface - whether through habitat encroachment or due to wildlife adapting to urban environments - disrupt species interactions and ecosystem processes in a variety of ways. They alter host-pathogen dynamics, reduce viable wildlife habitat, and lead to genetic isolation and physiological stress in animal populations.^3-6^ Moreover, closer proximity between domestic and wild animals further facilitates disease spillover, leading to infection of naïve species and introduction of pathogens into new ecosystems.^1,7^

Of the more than 200 known zoonotic diseases, approximately 60% originate from animals, contributing significantly to the global infectious disease burden.^8^ Studies have identified 56 zoonoses that are responsible for 2.7 million human deaths annually.^9^ Although wild and domestic animals are often perceived as sources of disease risks to humans needing to be controlled, animals can also be exploited as sentinels for zoonotic disease surveillance.^10^ Animal disease surveillance also has considerable value for conservation, given the rise in disease-associated mass mortalities in wild animals^11^ and the role humans can play in transmitting pathogens to other species, as seen with SARS-CoV-2 spillover into household pets (*e*.*g*., cats), zoo and farmed animals (*e*.*g*., tigers and minks, respectively) and wildlife (*e*.*g*., deer).^11-16^

Despite the acknowledged value of animal disease surveillance, effectively and sustainably implementing surveillance programs is difficult, especially for wildlife. One of the primary logistical challenges that has hindered these surveillance efforts is sample acquisition.^17,18^ Detecting the presence of pathogens in wildlife populations requires the collection of biological samples^19^ but, traditional sampling methods often require the capture and immobilization of wild animals, which pose significant risks to both animals and researchers.^20-27^ Besides the potential risk of animal injury and mortality related to the capture, capture-related stress can have long-term consequences for animals, including changes in behavior, losses in body condition, and lower reproductive success.^27-33^ While non-invasive sampling techniques - such as collecting feces, hair, urine, or saliva naturally shed or deposited into the environment - may mitigate these risks, they often require that researchers are present at the time of sample collection impeding sample collection across large, remote landscapes.^21,26,34-36^ New technologies are crucial for overcoming these limitations.^37^ Here, we describe the development of an innovative non-invasive device for animal disease sampling. Our device consists of a disposable plastic cassette that utilizes odor attractants to lure the animal into licking a filter paper that can be used for downstream pathogen analysis. To validate this device, first, we tested whether filter paper was an appropriate medium for long-term preservation of saliva-derived pathogen material. Second, we performed a series of animal trials evaluating the detectability of pathogen RNA in samples obtained from the device using RTqPCR. Finally, we used a high-throughput sequencing approach to explore the range of pathogen organisms detectable with our device. Our results indicate the potential value of this novel approach to disease monitoring in animals.

## Results

### Sampling device

Our sampler consists of a disposable 3D-printed cassette containing an attractant odor hidden behind a WhatmanTM™ filter paper circle that lures the target animal into licking a Whatman™ filter paper through the use of odor attractants (Figure 1A). The act of licking the filter paper results in the deposition of biological fluid (saliva) onto the paper, allowing for preservation and retrieval of genetic/biochemical material (Figure 1B-C). Innocuous food odors were tested to determine the best attractants for each target species in our study. For mice ethyl butyrate (“tutti-frutti” odor) adsorbed in ethylene-vinyl acetate beads was the best attractor. For cats, treat paste Churu™ (Inaba) was the best attractor.

**Figure 1.**
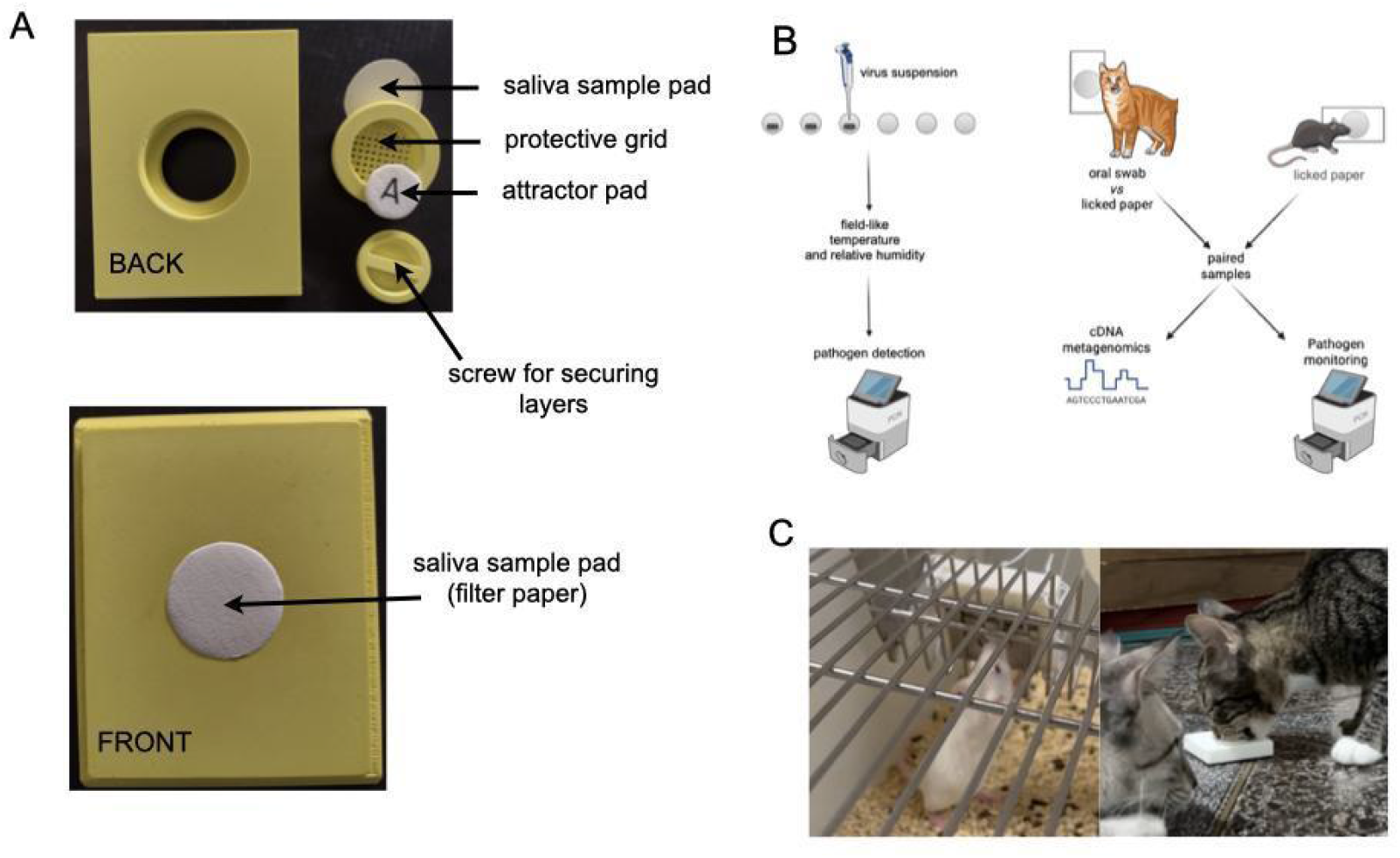
(A) Main features of single-use collection pad. The filter paper containing the attractor is secured behind the saliva collecting filter paper, with a protective grid in between. (B) Experimental design of this work. (C) Saliva collector being licked by laboratory mouse and shelter cat.

### Filter paper is a viable preservation medium for saliva-based virus detection

To evaluate the feasibility of our saliva-based molecular surveillance approach, we characterized rabies virus (RABV) RNA, zika virus (ZIKV) RNA, and herpesvirus simplex 1 and 2 DNA (HSV1, HSV2) stability in spiked saliva samples stored for different amounts of time (0, 3, 7, 14, or 30 days), under different temperature conditions (low humidity and hot [LHH], high humidity and hot [HHH}, room temperature [RT]), and on different types of filter paper (Whatman 3 or 54). On each day of the study, replicate samples stored under these different conditions were subject to RTqPCR to quantify virus recovery. RABV stability was assessed using only a single paper type (Whatman 3), and in a model testing the effects of day, temperature treatment, and their interaction on viral recovery, we found that only day emerged as a significant factor (Table S1). Surprisingly, RABV virus Cq values increased non-linearly with time (Figure 2A). For HSV1, HSV2, and ZIKV, our models tested the effects of day, temperature treatment, paper type, and all possible 2- and 3-way interactions on virus recovery. For HSV1, none of these factors affected virus stability (Table S1). For HSV2 and ZIKV, day interacted with temperature treatment to drive significant variation in virus stability (Table S1). In both cases, virus Cq values increased with time in the HHH treatment, but remained stable in the LHH and RT treatments (HSV2: Figure 2B, Table S2; ZIKV: Figure 2C, Table S3).

**Figure 2.**
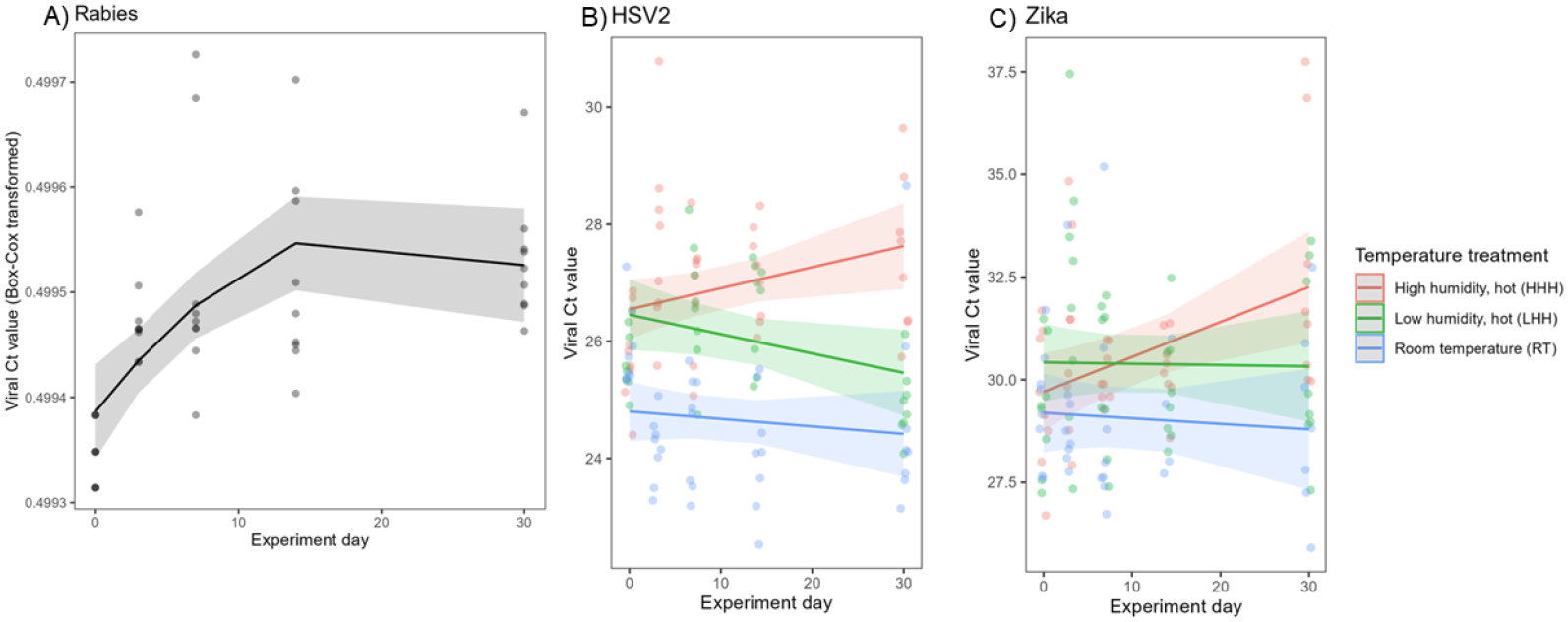
Predictors of variation in viral Cq values. A) Rabies variation was explained by day, with viral Cq increasing non-lineary with day; B) HSV2 and C) Zika values were explained by an interaction between temperature treatment and day, with Cq value increasing with day only under high humidity hot (HHH) conditions.

### RTqPCR detects host genes and virus in saliva samples obtained from the sampling device

To test our ability to detect viruses from saliva collected with the sampling device, we used mice experimentally infected with RABV and shelter cats naturally infected with Feline Leukemia Virus (FeLV) and/or Feline Immunodeficiency virus (FIV). In mice, we performed two distinct experiments using the AgV3 (experiment 1) and CVS (experiment 2) strains of RABV. For each experiment, 10 mice were given 10 μL of a 10^6.5^ LD_50%_/mL RABV suspension per nostril and housed together in a single cage. The saliva sampler was installed on the external side of the metal grid cover of each cage (Figure 1C). The sample collector was installed two days before infection, and filter paper was changed daily. To verify mouse contact with the sampler, we screened for murine beta-actin RNA using RTqPCR. We consistently detected murine beta-actin for the duration of both experiments, however, detection levels varied significantly over time for both experiments (experiment 1: F = 140.3, df = 8, p <0.001, Figure 3A; experiment 2: F = 166.1, df = 10, p <0.001, Figure 3B);. Although this assay is not saliva specific, it confirms daily host contact with the filter paper. For both RABV strains, mice presented clinical signs of infection and were dead by day 10 (experiment 1) and 10-13 of infection (experiment 2) (*survival curves not shown*), but no virus was detected in collected saliva samples.

**Figure 3.**
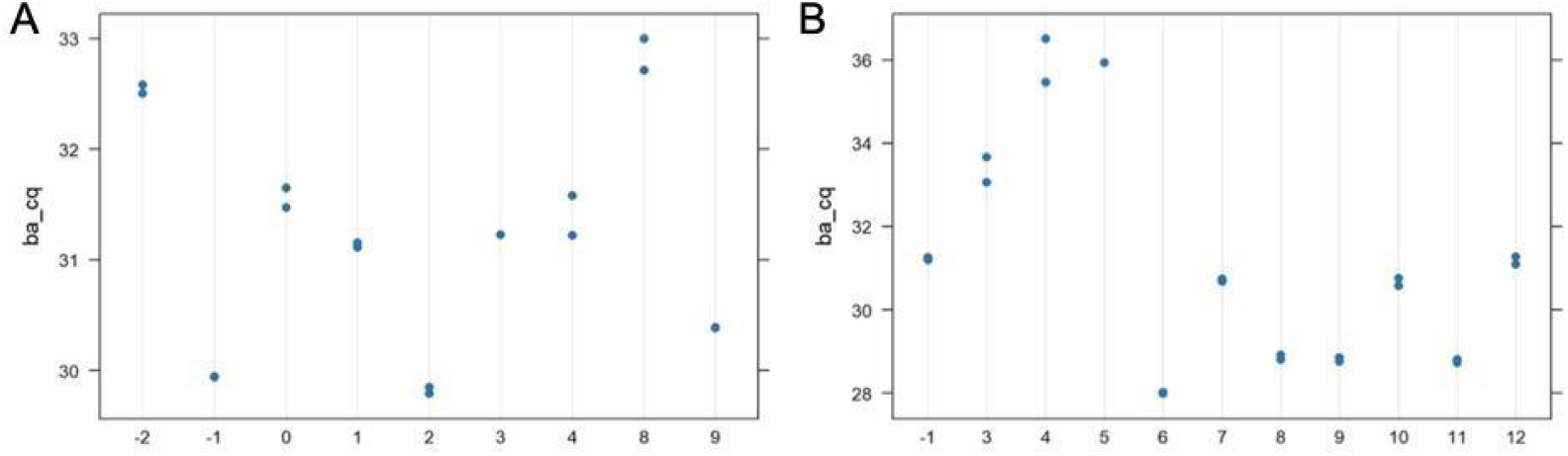
Variation in beta-actin gene Cq values from licked paper obtained from collector devices installed in mouse cages during experimental rabies infection with the A) AgV3 and B) CVS RABV strains. In both cases, beta-actin was detected on each day of the experiment, but levels of detection varied across days. Samples were run in duplicate on each day.

We used domestic cats naturally infected with FeLV and/or FIV to further test the efficacy of the sampling device. Forty individual cats of both sexes, multiple ages, and breeds housed in a rescue and rehabilitation center were used. Animals were housed singly, or in groups of three for family groups, in dens of 1.2 m^2^ of floor area and 3 m of height, with water and food containers provided *ad libitum* (Figure S1). Upon admittance to the facility all animals were screened for FIV and FeLV using the Idexx SNAP™ serological test. A single sample collector was introduced into each den for 10 minutes (Figure 1C), and interactions between animals and the sampling device (*e*.*g*. licking, nose touching, sneezing) were monitored using video recordings and direct observation. After the sampler was retrieved, each animal in the den received an oral swab. Samples were screened for FeLV and FIV RNA by RTqPCR. Sampling was performed on a monthly basis for 6 rounds; some animals were sampled more than once due to their extended stay at the facility.

Only five individuals tested seropositive for FIV across the study, only two of which tested RTqPCR positive, so we focused our formal analyses on FeLV. For analysis, we included only animals whose RTqPCR results had duplicate concordance and correct melt peaks and excluded samples with ambiguous results. Whenever a sample was negative for both FeLV and FIV, the sample was amplified for *F. catus* 28S to ensure sample adequacy. Thirteen individuals were seropositive for FeLV upon admittance to the facility, while 18 returned RTqPCR positive swabs and 12 returned RTqPCR positive licked papers over the course of the study. Both the swab (binomial GLMM; n = 43 samples, 29 unique cats: estimate = 20.75, z = 3.761, = 0.000169) and paper (n = 47 samples, 35 unique cats: estimate = 24.08, z = 3.736, p = 0.000187) methods significantly predicted serological status. We used a subset of 37 samples that generated viable RTqPCR swab and paper results to test for agreement between the two methods and found moderate agreement (kappa = 0.409, z = 2.52, p = 0.0117). Finally, when comparing the Cq values for the subset of 13 samples for which both the swab and licked paper returned positive tests, we found that FeLV Cq values were significantly higher for the paper samples (Wilcoxon test: W = 91, p = 0.0002, Figure 4).

**Figure 4.**
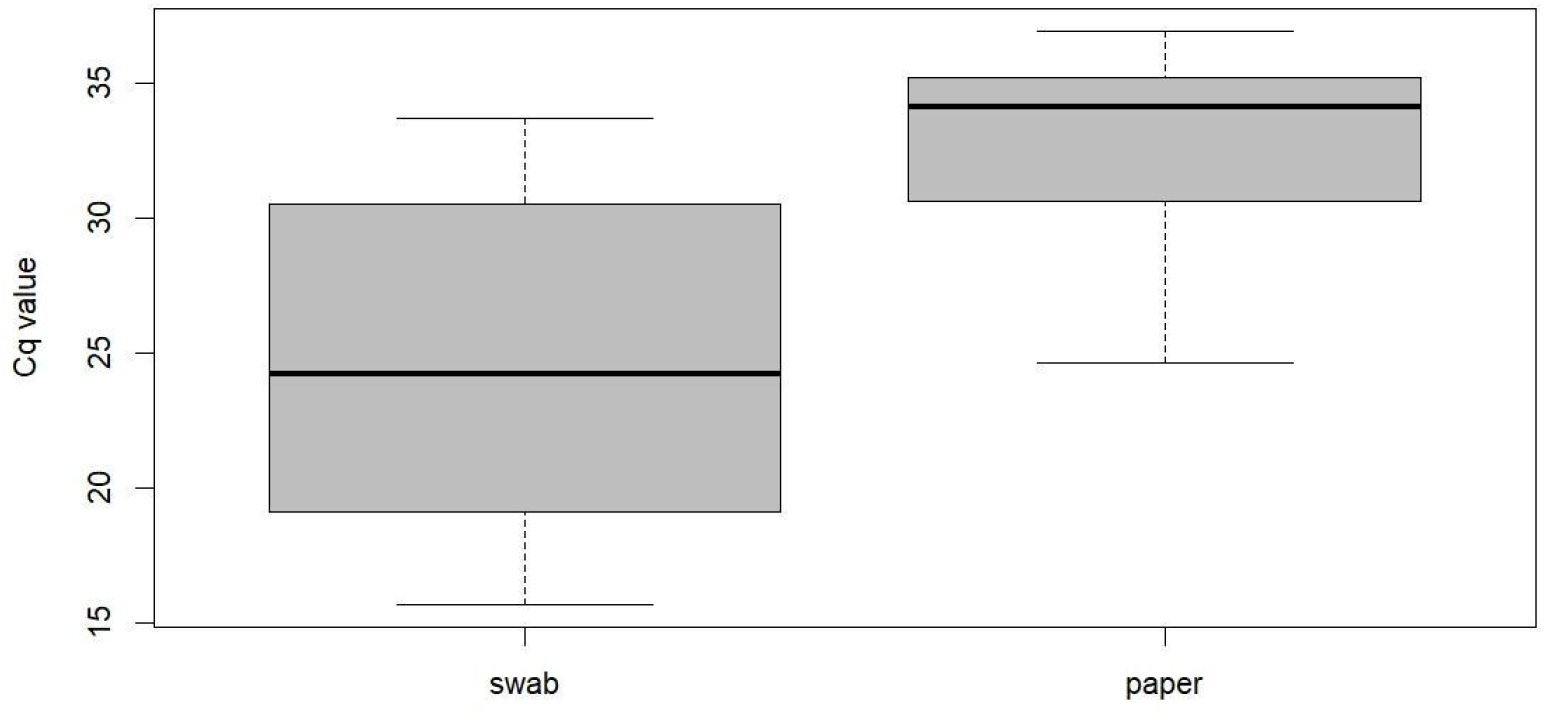
Comparison of FeLV paired Cq values from oral swabs and licked paper for 13 cats. Viral Cq values were significantly higher for the paper samples.

### High-throughput sequencing of sampling device-derived saliva detects a wide range of pathogens and symbionts

To characterize a broader set of pathogens detectable in device-derived saliva samples, we subjected a subset of samples we collected from mice and cats to long-read cDNA metabarcoding. Annotation of 4.86M reads identified a myriad of bacteria, archaea, fungi, plants, and eukaryotic species present in animal saliva samples (Figures 5 and 6). Similar to the RTqPCR results, we did not detect RABV in the samples from experimentally infected mice. Figure 5 shows variation on the relative expression of mice-specific pathogens (upper panel) and beneficial symbionts (middle panel) during RABV infection, while chow components remained stable through the same period (lower panel). For the naturally infected cats, FIV was not detected in any sample by sequencing (n = 32), including samples (n = 5) that tested positive for serology.

**Figure 5.**
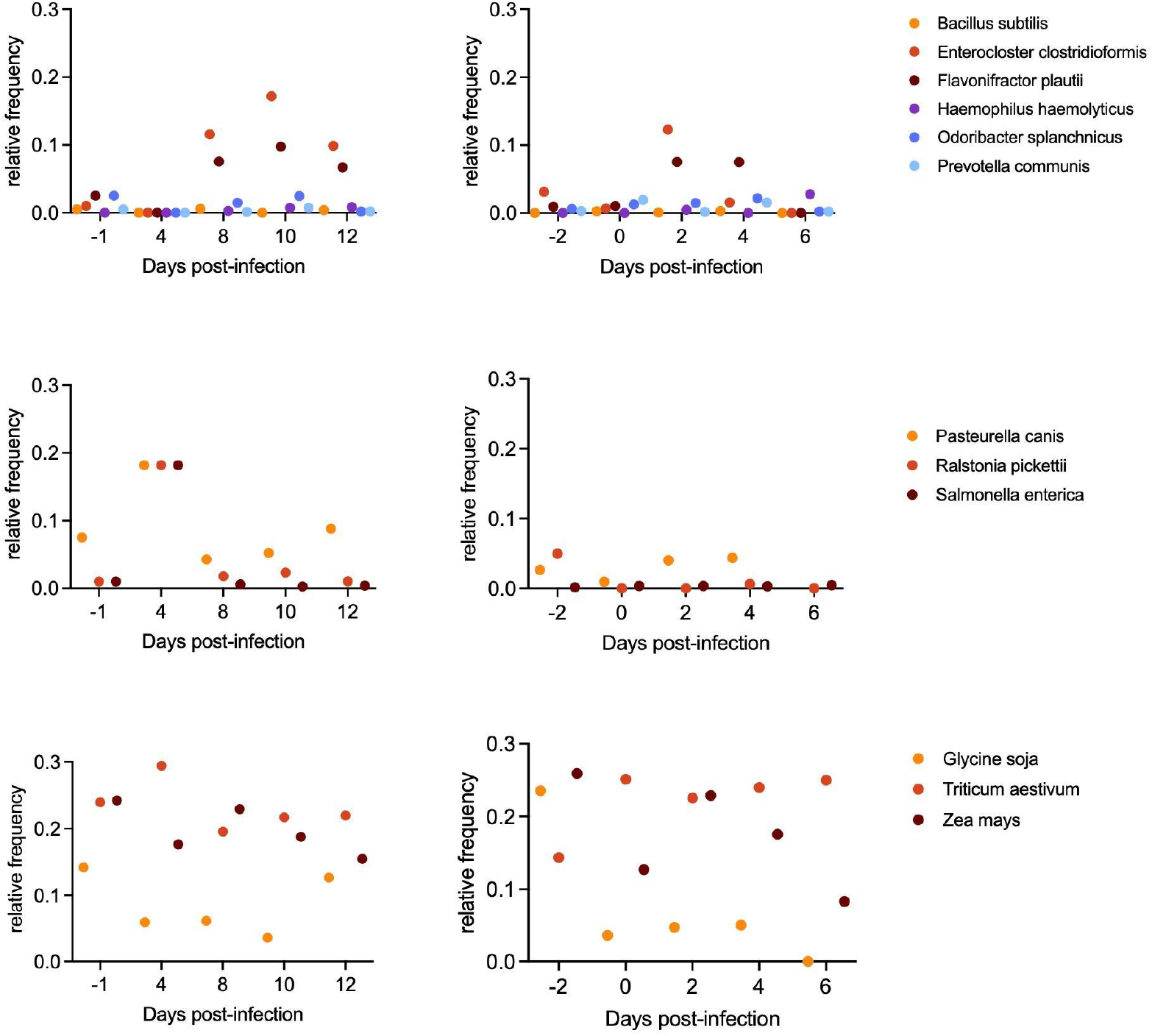
Taxonomic composition of mice saliva samples over the course of rabies infection. Proportion of each most frequent species found across mice samples based on the hits detected in each sample. (Upper panels) Beneficial bacteria of the mouse microbiome. (Middle panels) Mice-specific pathogens. (Lower panels) Chow components. All values are represented as 95% CI (Wilson method).

**Figure 6.**
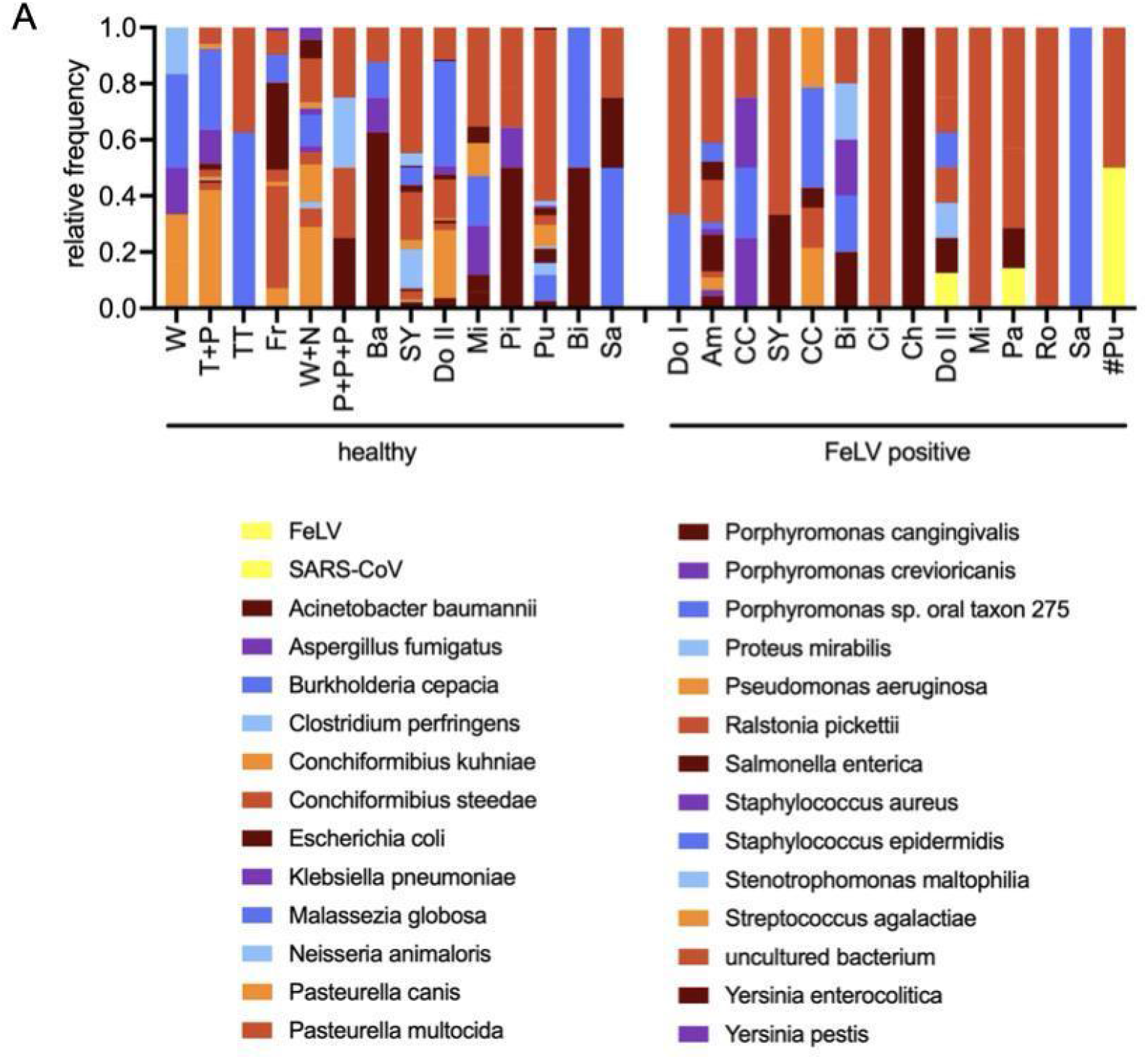
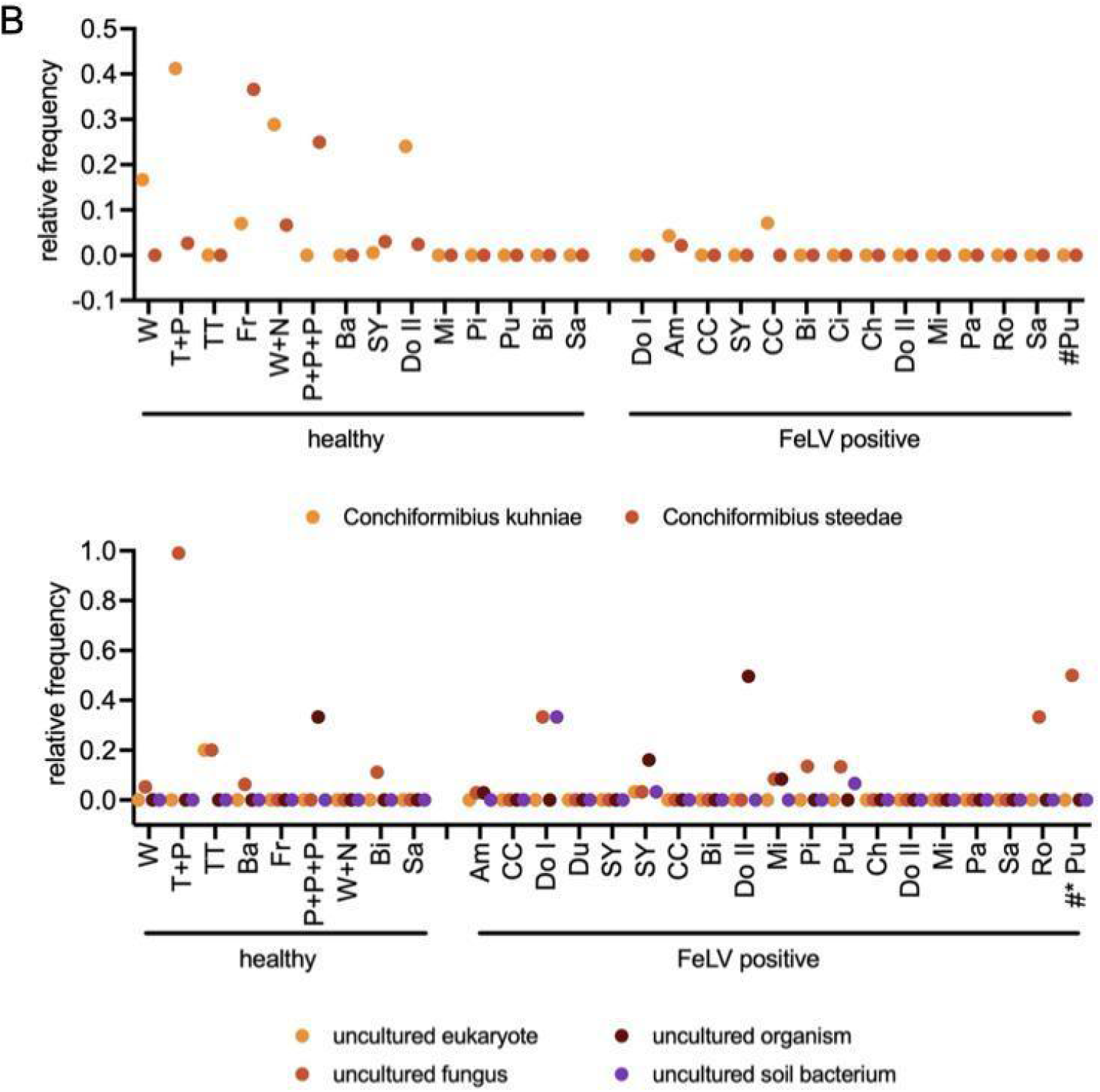
Taxonomic composition of cats saliva samples. Proportion of each most frequent species found across cats samples based on the hits detected in each sample. All values are represented as 95% CI (Wilson method). A) Feline-specific pathogens. The symbol # indicates a SARS-CoV-positive cat. B) Relative frequency of *Conchiformibius kuhniae* and *C. steedae*, beneficial bacteria of the cat’s oral microbiome (upper panel) and of uncultured organisms (lower panel).

However, FeLV was detected in three samples that were positive for FeLV on serology and RTqPCR of licked paper (Table S4). We also detected SARS-CoV-2 in one sample (Figure 6A and Table S4). As expected for individual cats coming from different areas and health/nutritional conditions, the taxonomic composition of saliva was heterogeneous (Figure 6A). We found a higher relative frequency of beneficial symbionts in healthy cats, while unidentified organisms were detected in several samples (Figure 6B). Note that samples from some cats initially identified as serologically negative for FeLV did not show the beneficial symbionts. Posterior samples from the same cats revealed they had become positive for FeLV and continued not presenting the symbionts (Figure 6B).

## Discussion

The sampling methodology reported here provides a capture-free means to study environmental nucleic-acid signatures via mammals’ exploratory behavior. As we have shown for FeLV in cats, it is possible to obtain insights about infection status of individual animals by using cost-effective sampling and detection assays (RTqPCR). The negative impact the capture has on wildlife welfare and survival limits our ability to study endangered species. Our methodology can be used to study these species since the saliva collection occurs in a completely voluntary manner and does not require the presence of the researcher. Our methodology is likely to be of great utility for monitoring pathogen circulation in a given environment, and for guiding proactive mitigation measures before outbreaks occur.

FIV virus load can be low and notoriously difficult to be caught in circulation, thus the preferred method of diagnosis is the detection of antibodies against FIV on SNAP™ test (Idexx) instead of virus antigens.^38,39^ Accordingly, we did not find FIV in any sequenced samples of seropositive cats. We were also unable to detect RABV by either RTqPCR or sequencing of saliva samples. RABV shedding is intermittent, and intranasal infection, as used in our study, has the potential to directly infect CNS tissues, bypassing salivary gland infection.^40-43^ A more detailed study on the kinetics of both FIV and RABV infection and shedding in saliva is beyond the scope of our study. In fact, our assumption for the valuation of our methodology is not necessarily diagnosing the animal that interacts with the saliva collector, but rather using that animal’s saliva as a proxy to represent its environment. Mammals are natural explorers of their environment, and our methodology aims at uncovering what is circulating in a larger scale environment by examining what pathogens mammals have been exposed to. Our ongoing research has successfully collected saliva samples of Andean bears living in captivity and free-ranging capybaras, marmosets, opossums, and wild rodents.

We recognize that our protocol for sequencing needs fine-tuning to increase depth of sequencing and that as it is, we are exploring only the most expressed transcripts and RNA genomes rather than the total molecular content. Our detections represent nucleic-acid signatures captured from saliva on paper matrices, consistent with molecular surveillance approaches where viability is inferred and strengthened by complementary assays. Nevertheless, our sequencing results reveal the richness of the gene space, demonstrating the potential of our methodology for discovering rare or unknown species that circulate in the environment of the saliva donor, enabling this player to be recognized as an environmental sentinel.

As a demonstration for this point, SARS-CoV-2 was identified by sequencing the saliva sample of one cat with respiratory tract disease. SARS-CoV-2 is not routinely monitored in domestic cats, despite the fact that it does infect these animals that might serve as potential spreaders to humans and other species. This finding highlights the potential of metagenomics sequencing of saliva samples as a tool for unbiased monitoring at landscape scale. Interpretation of our findings as infectious viable pathogens will require viability assays.

A major limitation of our study is that our metabarcoding sequencing was based on polyadenylated RNA, minute amounts of starting material and relatively shallow coverage of the species found by sequencing. Future users of this methodology can easily adapt their pipeline for DNA sequencing, which should provide larger material amounts and deeper coverage. We also did not use custom databases for raw data analysis, instead we used user-friendly freely available tools (CZID.org). The use of custom-made databases focusing on specific targets of interest might also improve coverage of the sequencing. Nevertheless, our results support microbial RNA recovery from saliva that can be used for environmental monitoring and discovery.

## Conclusions

In controlled settings, a 3-D printed cassette with odor attractants elicited voluntary licking and saliva deposition on filter paper by cats and mice, allowing for detection of RNA viruses by RTqPCR and unbiased microbial and environmental discovery by third-generation sequencing. This study demonstrates feasibility in sheltered/captive deployments, providing necessary foundations for field calibration.

## Methods

### Ethical considerations

Sampling of the shelter cats was only conducted after informed consent had been signed by the shelter staff for each participating cat, and upon protocol approval by CEUA-ICB (Institutional Committee for Animal Use and Care, ICB-USP, 6722160123). Sampling of the mice was approved by CEUA-ICB (6722160123) and RABV infection was approved by the CEUA-FMVZ (7776070623).

### Spontaneous Saliva Samples Collection

Our sampler consists of a disposable 3D-printed cassette containing an attractant odor hidden behind a WhatmanTM™ filter paper circle that is licked by the target animal, depositing oral-contact biological material onto the paper, and allowing for preservation and retrieval of genetic/biochemical material (Figure 1B). Innocuous food odors were tested to determine best attractants for each target species. Ethyl Butyrate (“tutti-frutti” odor) adsorbed in ethylene-vinyl acetate beads (both kind gifts by Givaudan) was the best attractor for mice, while treat paste Churu™ (Inaba) on a piece of filter paper was the best bait for cats.

### Viral Decay in Saliva Stored on Paper

Human saliva pools (5 donors) were spiked with known amounts of tissue culture-grown pathogens and three *μ*L were added to four mm filter paper punches, stored under the following conditions: room temperature [RT] (23°C, relative humidity (RH) ∼65%), low humidity and hot [LHH] (45°C, RH *≤*16%), and high humidity and hot [HHH} (45°C, RH ∼85%). All conditions were monitored and controlled using thermometers and hygrometers. Triplicate punches were processed on days 0 (fresh), 1, 7, 14, 30, and 60, or otherwise noted, using Instagene (Bio-Rad) and Proteinase K (Thermofisher) as detailed below. Pathogen DNA/RNA was quantified by (RT)qPCR using primers broadly validated in the literature and cycling conditions according to manufacturer’s instructions of Power SYBR® RNA-to-CT™ 1-Step Kit (ThermoFisher). Zika virus (ZIKV) primers were ZIKV-forward - 5’-CCGCTGCCCAACACAAG-3’ and ZIKV-reverse - 5’-CCACTAACGTTCTTTTGCAGACAT-3’.^53^ Herpes Simplex Virus 1 (HSV1) primers were HSV1-forward - 5’-GTTGAGCTAGCCAGCGA-3’ and HSV1-reverse- 5’-CGTTAAGGACCTTGGTGAGC-3’.^54^ Herpes Simplex Virus 2 (HSV2) primers were HSV2-forward - 5’-CACACCACACGACAACAA-3’ and HSV2-reverse - 5’-TAGTTCAAACACGGAAGCC-3’. ^54^ Reactions for Rabies virus (RABV) used the primers listed below. Positive controls were DNA/RNA extracted from fresh tissue cultured pathogens.

### 3D Printing of Saliva Collectors

Collectors were printed using PLA filament using an Enders 3SE printer (both by Creality). STL files for 3D printing the collectors are available through the link: https://osf.io/cr68g/?view_only=21f5ab7be2574d6795cf66f40edc1c5a

### Experimental infection of mice with Rabies virus (RABV)

The CVS (Challenge Virus Standard)-1 strain or the AgV3 strain of RABV with a titer of 10^6.5^ LD_50%_/mL cultured in BHK-21 (baby hamster kidney) were used for the experimental infections. Isogenic mice of the Balb/c lineage (n = 20) were purchased from ANILAB Com. Ltda. (Brazil), at 14 days of age, and kept at the School of Veterinary Medicine (FMVZ-USP) Animal Facility with water and food *ad libitum*, and a 12h light period throughout their stay, including the experimental periods. For the RABV inoculation, mice were anesthetized with Isoflurane and given 10 μL of RABV suspension per nostril. Mice were caged in groups of 10 and the saliva sampler was installed on the external side of the metal grid cover of the cage, behind the metal hole that holds one of the water bottles (Figure 1C). The sample collector was installed two days before infection, and the filter paper was exchanged daily. All steps were carried out at the Rabies Laboratory, Department of Preventive Veterinary Medicine and Animal Health (FMVZ-USP), Brazil.

### Cats Naturally Infected with Feline Immunodeficiency Virus (FIV) and/or Feline Leukemia Virus (FeLV)

Sixty domestic cats, both sexes, all ages, and breeds participated in sampling. They lived in a non-governmental organization for cat rescue and rehabilitation (“Enquanto Houver Chance”, São Paulo, Brazil), kept in individual dens of 1.2 m^2^ of floor area and 3 m of height. The back wall was tiled and three sides were fenced. They had individual water and food containers, provided *ad libitum*, and individual litter boxes. The back wall had several climbing boards and wood boxes as environmental enrichment (Figure S1). Thirty cats were FeLV-positive and 10 were FIV-positive based on serological test (Idexx SNAP test™) at time of shelter admittance, and the others tested negative. Sampling was conducted on a monthly basis for 6 rounds. Some animals were sampled more than one time, pending on their extended stay at the shelter. A sample collector was left for 10 minutes in the presence of one individual cat or up to three cats whenever they were a family housed together. After 10 minutes, the collector was retrieved, an oral swab performed, and the samples were carried back to the laboratory in an ice box. All licked papers and swabs were kept at −80°C until processing. Power analysis for FeLV, considering Cq values obtained on RTqPCR, resulted in achieved power = 0.843, alpha = 0.05 (two-tailed), considering 16 animals (8 positives and 8 negatives).

### Manual Saliva Samples Collection in Cats

For validation of the sampling device as an efficient collection method, approximately ⪝20 *μ*L of saliva wascollected via oral swabbing of the cats after a positive interaction with the saliva collector. The swabs were carried back to the laboratory in an ice box and stored at −80°C until tested. In cases where members of the same family were housed together, we had a sample collector licked by all the animals (“collective paper”) but individually collected oral swabs.

### Samples processing

Seven punches of 4 mm were recovered from the licked papers or four pieces (∼4 mm long) were cut out of the swabs and placed into microtubes containing 600 μL of Lysis buffer (PureLink RNA Mini kit, Thermofisher) and proceeded according to the manufacturer’s protocol. Alternatively, viral decayment experiment punches and mice samples were processed adding 200 *μ*L of Instagene (Bio-Rad) and 14 *μ*L of proteinase K [10mg/mL] to paper punches in triplicate and incubating them at 56°C for 30 min, followed by vortexing for 10 secs. The mixture was heated at 100°C for 8 min, vortexed for 10 secs, and centrifuged at 12,000 *xg* for 3 min. The supernatant was used for downstream applications. Punches of paper that were exposed to the environment but not licked by the animals were extracted as water control. RNAs were stored at −80°C until use. In the case of Instagene-extracted samples, upon defrosting samples were vortexed for 10 secs and centrifuged at 12,000 *xg* for 3 min. For sequencing run 2, RNA degradation was evaluated by High Sensitivity Tapestation (Agilent) (Figure S2).

### RABV Confirmation of Infection by RTqPCR

Total RNA was extracted from mice brains immediately upon death using Instagene (Bio-Rad). SYBR green RTqPCR targeting the RABV N gene, using the Beta-actin gene as an internal reference, was used to assess the pathogen load by comparison against a titration curve of a known number of copies of the RABV gene, as described in.^44^ RTqPCRs were performed using primers JW12 - 5’-ATGTAACACCYCTACAATG-3’ and primer N165-148 5’-GCAGGGTAYTTRTACTCATA-3’ (all reagents from Promega). Positive controls were RNA extracted from BHK-21 cell cultures infected with RABV strain CVS. Murine ϐ-actin was used as internal control.

### FeLV / FIV Confirmation of Infection by RTqPCR

FeLV exoU3 and FIV 5’UTR genes were quantified by RTqPCR using primers broadly validated in the literature. FeLV primers were FeLV-exoU3-forward - 5’-AACAGCAGAAGTTTCAAGGCC-3’ and FeLV-exoU3-reverse - 5’-TTATAGCAGAAAGCGCGCG-3’(2005 TANDOM).^45^ FIV primers were FIV_gag_upstr - 5’-ATGGGGAAYGGACAGGGGCGAGA-3’ and FIV_gag_downstr - 5’-TCTGGTATRTCACCAGGTTCTCGTCCTGTA-3’.^39^ As internal control, Fcatus_28S_fwd - 5’-CGCTAATAGGGAATGTGAGCTAGG-3’ and Fcatus_28S_rev - 5’- TGTCTGAACCTCCAGTTTCTCTGG-3’.^46^ Reagents and cycling conditions of Power SYBR® RNA-to-CT™ 1-Step Kit (Thermofisher). Positive controls were DNA extracted from clinical samples from FeLV-positive cats and plasmids containing FIV 5’UTR.

### Library Preparation and High-throughput Sequencing

For cDNA metabarcoding, purified total RNA and controls (extraction control samples (water), positive controls (FeLV, FIV, and RABV), and barcoding control from kit) were polyadenylated for 1 minute at 37°C using 5U of *E. coli* Poly(A) Polymerase (New England Biolabs, NEB), followed by cleaning with RNAclean XP beads (Beckman Coulter) (according to ONT protocol). Polyadenylated RNA was ligated to a double stranded adapter (Oxford Nanopore Technology, ONT) using NEBNext Quick Ligation Reaction Buffer, T4 DNA Ligase (NEB), and RNase OUT (Thermofisher) and after adapter digestion with Lambda Exonulease and USER (both by NEB), cleaned-up with RNAClean XP beads (Beckman Coulter) using the Short Fragment Buffer (ONT) for washes. Reverse transcription and strand-switching were performed using Maxima H Minus Reverse Transcriptase (Thermofisher), RNase OUT (ThermoFisher) and Strand Switching Primer II (from the cDNA-PCR Sequencing V14 Barcoding kit, SQK-PCB114.24, ONT).

Double-stranded DNAs were barcoded using unique primers from SQK-PCB114.24 (ONT) for combined sequencing. All samples from Run 1 were subjected to an extension cycle of 1 min at 65°C for barcoding, while samples from Run 2 (except the positive control) were subjected to 20 extension cycles of 2 mins at 65°C. The positive control (75ng of *S. ceverisiae* enolase RNA (RCS-001 from ONT)) was subjected to a single 1 min extension cycle at 65°C. Barcoded samples were cleaned with AMPure XP beads (Beckman Coulter), pooled (10 samples from mice and 10 samples from cats in sequencing run 1, and 22 samples from cats in sequencing run 2) and sequencing adaptors (ONT) were ligated. Pooling of the 1st library considered equal volumes of samples, regardless of cDNA concentration. Pooling of the 2nd library considered Tapestation results (Figure S2) to each near equimolar concentrations. Libraries were loaded onto a MinION flow cell (R10 version) and run for 16h30m (sequencing run 1) or 69h40m (sequencing run 2) on a MinION 1kB sequencer (ONT). Run 1 yielded 1.19M reads with score Q > 8, while the run 2 yielded 3.67 M reads with score Q > 9. Figure S3 shows the pore occupancy reports from both runs. Pore occupancy in the first run was below ideal but improved in the second run by the addition of RCS to the library, and as a result yielded longer reads.

### Virus and Other Taxon Detection Workflow

Basecalling and demultiplexing were performed with the MinKNOW operating software (ONT).^47^ Raw reads were trimmed to remove adapter sequences and then filtered to remove reads with q-scores ≤ 9 and read lengths ≤ 100 bp.^48,49^ The processed reads were aligned to the NCBI RefSeq database and visualized using CZID.^50^ Taxon-based read counts were obtained from CZID with default settings optimized for nanopore data.^50^

### Statistical Analyses

For the RTqPCR assays, we used Thermofisher Cloud to analyze and export Cq values. To test for effects of day, temperature treatment, filter paper type, and all possible interactions on virus stability (quantified at Cq value), we fit separate linear mixed effect models (LMM) and then used backwards stepwise elimination to determine which of these predictors contributed to significant variation in virus Cq values. Because pieces of filter paper were sampled in triplicate, all models also included replicate identity as a random effect. To achieve normally distributed model residuals, rabies virus Cq data was Box Cox transformed and a quadratic polynomial term for day was added to the full model. The Zika virus Cq data had a single influential outlier which significantly altered the results of the stepwise elimination and downstream analyses, so we removed this zero value (which was unique across the entire dataset) from the analyses presented in the main text. Results including the zero outlier are provided in the supplement (see Table S5-S6). All analyses were performed in R (version 4.4.1).51 LMMs were implemented using the lme4 and lmerTest packages with backward elimination performed with the *step* function. Post hoc analyses were performed using emmeans and model checks were implemented using DHARMa.

To test whether the detection of murine beta actin varied over experimental days, we used ANOVA tests. To test whether RTqPCR results from oral swabs and licked paper were predictive of FeLV serological test results in cats, we used binomial generalized linear mixed models. The result of a one time serological test for each cat was included as the response variable and either the swab or paper RTqPCR test result (positive or negative) was included as a predictor. Because some cats were RTqPCR sampled more than once, we included cat identity as a random effect. Also, because some cats were housed in family groups that all had access to the same paper, we included the sampling event as a random effect in the paper analysis to account for the fact that multiple cats were exposed to the same paper in some instances. We then used Cohen’s Kappa to test for agreement between the swab and paper methods. Finally, we used Wilcoxon signed rank tests to compare FeLV Cq values for paired swab and paper samples. All analyses were performed in R (version 4.4.1).^51^ GLMMs were implemented using the lme4 package and the Kappa test was implemented in the irr package.

## Code availability

The code for the viral decay analyses and the STL files for 3D printing the collectors are available through the link: https://osf.io/cr68g/?view_only=21f5ab7be2574d6795cf66f40edc1c5a

## Data availability

Videos of the animals interacting with S.W.A.B. can be found at linktr.ee/SWABbiosampler

## Acknowledgements

We thank the financial support that made this study possible: Yale Planetary Solutions Seed Grant, Revive & Restore Catalyst Fund (2023-070), CNPq (National Council for Scientific and Technological Development, 309328/2022-5), University of São Paulo PUB Scholarship Program (to A.C.F.G.), CAPES scholarship (88887.928496 to D.N.H.B.), Wild Animal Initiative (SG 23-030), and FAPESP (São Paulo Research Foundation, 2022/07115-7). We thank Bridget Baumgartner for invaluable input in the early phases of this study and Mauricio Cella, from Givaudan for kindly providing the odors. We also acknowledge Candace Williams, Daniel Bruzzese, Daniel Lahr, Gabriel Gomide, João Marcelo Alves, Kushal Suryamohan, Mrinalini Watsa, and Renata Ferreira for invaluable advice on Nanopore sequencing, computing, and data analysis. We thank all the staff at the cat shelter and rehabilitation “Enquanto Houver Chance” for their good work with the cats and for facilitating our sample collection.

## Supplementary Material

**Figure S1.**
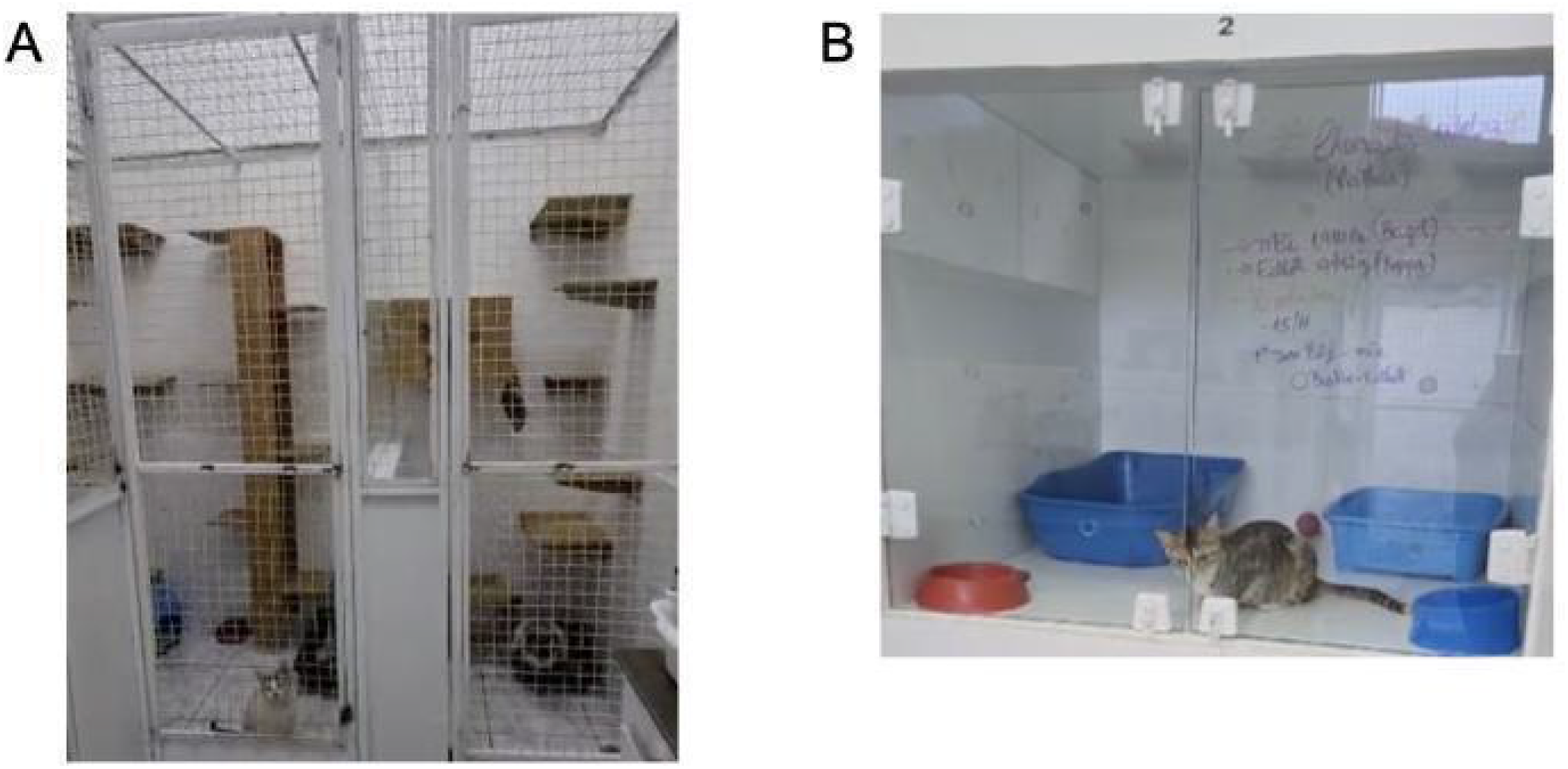
The participating shelter had a maximum occupancy of 40 cats in individual/family dens. A) Individual (right side) and family (left side) dens. B) Intensive care box.

**Figure S2.**
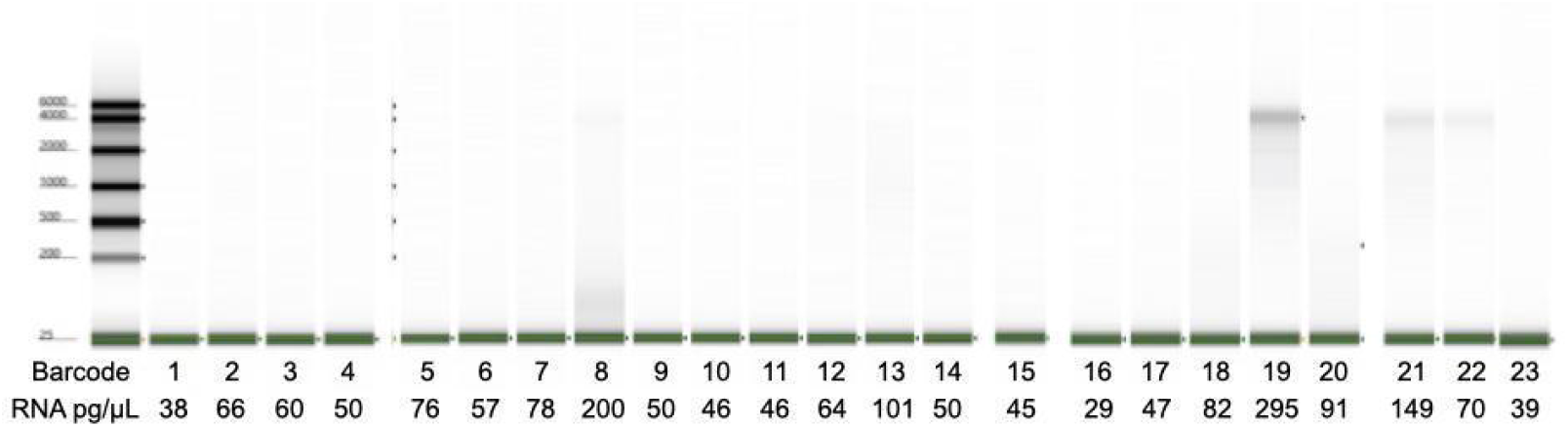
Tapestation results for cat RNA samples (sequencing run 2).

**Figure S3.**
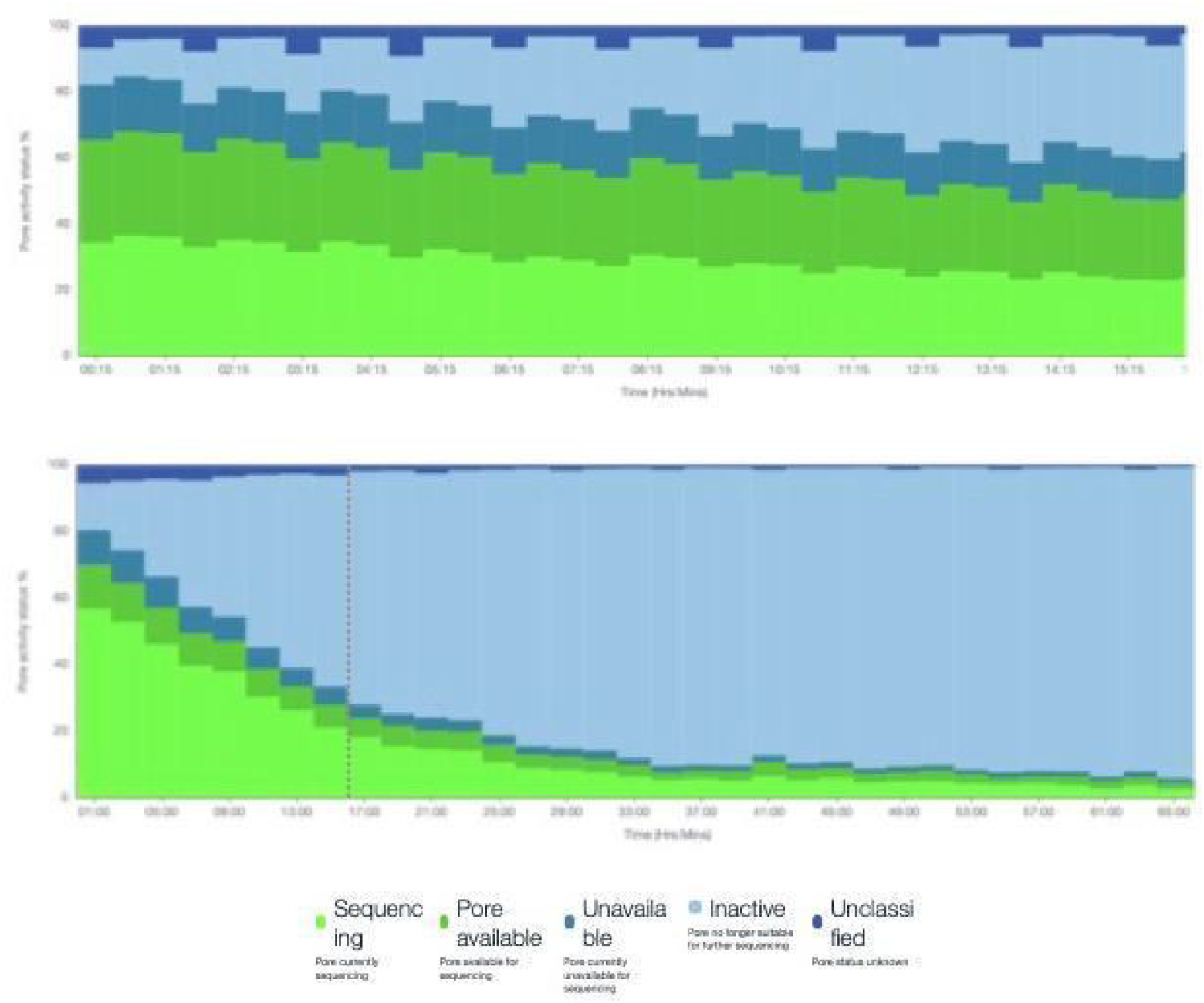
Nanopore MinION sequencing run reports. The vertical dotted grey line in the lower panel indicates 16hs (run 2) for comparison with the upper panel (run 1, total of 16hs of run).

**Table S1.** Results of linear mixed effects models with stepwise elimination examining predictors of variation in the Cq values of multiple viruses (Rabies, HSV1, HSV2, Zika). Terms included in the best fitting model are listed with significant terms in bold.

| Virus | (Intercept) | pape<br>r | temp | day | temp:day | poly(day, 2) |
| --- | --- | --- | --- | --- | --- | --- |
| Rabies | p = 2.13e-150 | - | - | - | - | **F = 11.94**<br>**p = 7.86e-05** |
| HSV1 | p = 4.39e-37 | - | - | - | - | - |
| HSV2 | p = 1.94e-108 | - | **F = 14.05**<br>**p = 3.88e-06** | F = 0.11<br>p = 0.745 | **F = 4.22**<br>**p = 0.0172** | - |
| Zika | p = 2.15e-58 | - | F = 1.66<br>p = 0.198 | F = 1.6<br>p = 0.21 | **F = 3.03**<br>**p = 0.0534** | - |

**Table S2.** Post hoc tests for HSV2 linear mixed effects model. Significant terms are listed in bold.

| temp | day.trend | SE | df | lower.CL | upper.CL | t.ratio | p.value |
| --- | --- | --- | --- | --- | --- | --- | --- |
| HHH | **0.036009** | **0.016834** | **85.21212** | **0.002539** | **0.069479** | **2.13902** | **0.035295** |
| LHH | -0.03301 | 0.018082 | 85.21212 | -0.06896 | 0.002943 | -1.82542 | 0.071442 |
| RT | -0.01276 | 0.016834 | 85.21212 | -0.04623 | 0.020706 | -0.75824 | 0.450398 |

**Table S3.**
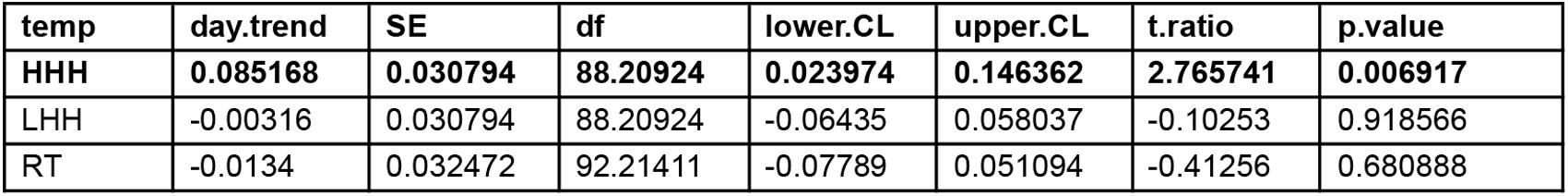
Post hoc tests for Zika virus linear mixed effects model. Significant terms are listed in bold.

| temp | day.trend | SE | df | lower.CL | upper.CL | t.ratio | p.value |
| --- | --- | --- | --- | --- | --- | --- | --- |
| HHH | **0.085168** | **0.030794** | **88.20924** | **0.023974** | **0.146362** | **2.765741** | **0.006917** |
| LHH | -0.00316 | 0.030794 | 88.20924 | -0.06435 | 0.058037 | -0.10253 | 0.918566 |
| RT | -0.0134 | 0.032472 | 92.21411 | -0.07789 | 0.051094 | -0.41256 | 0.680888 |

**Table S4.**
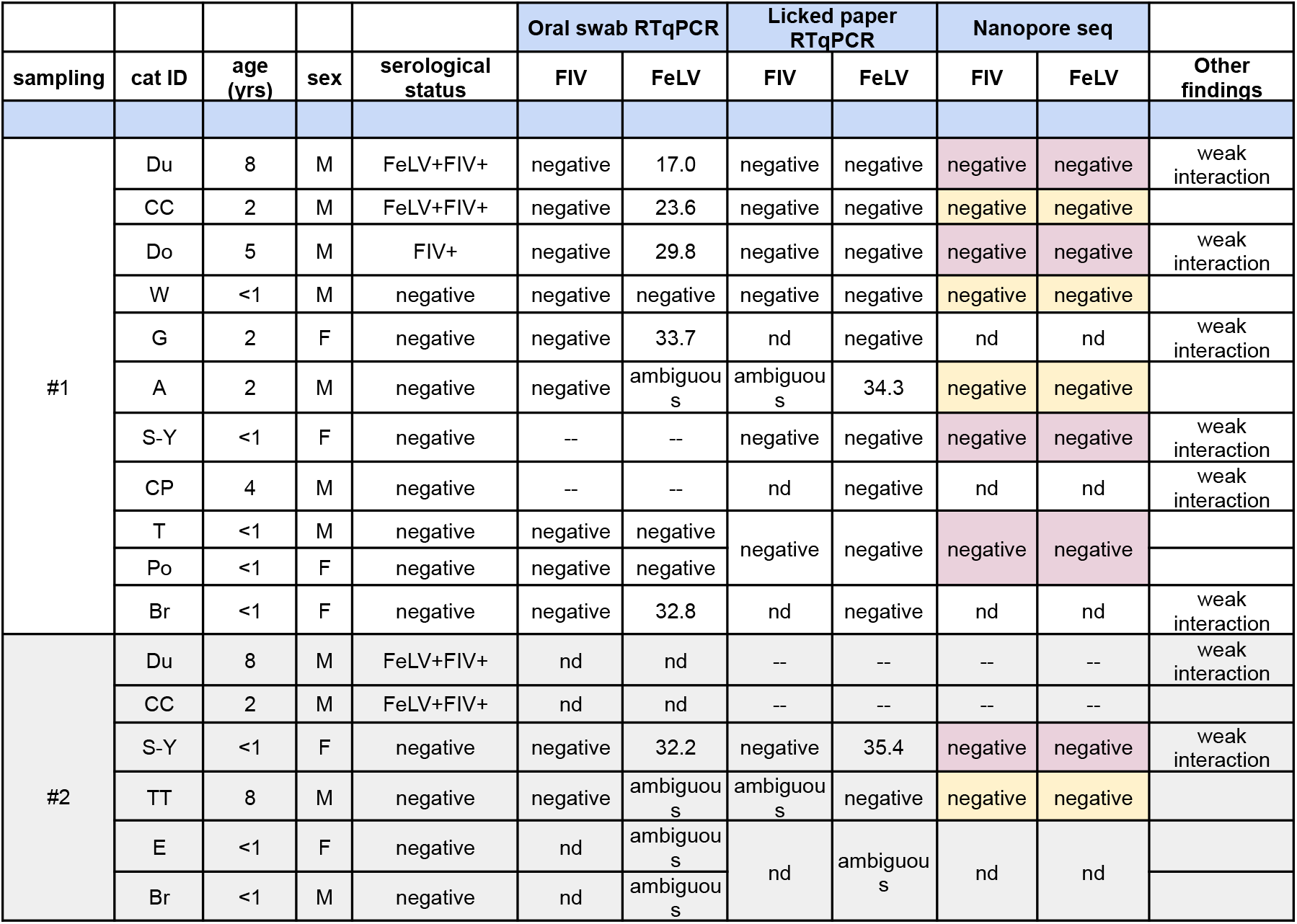
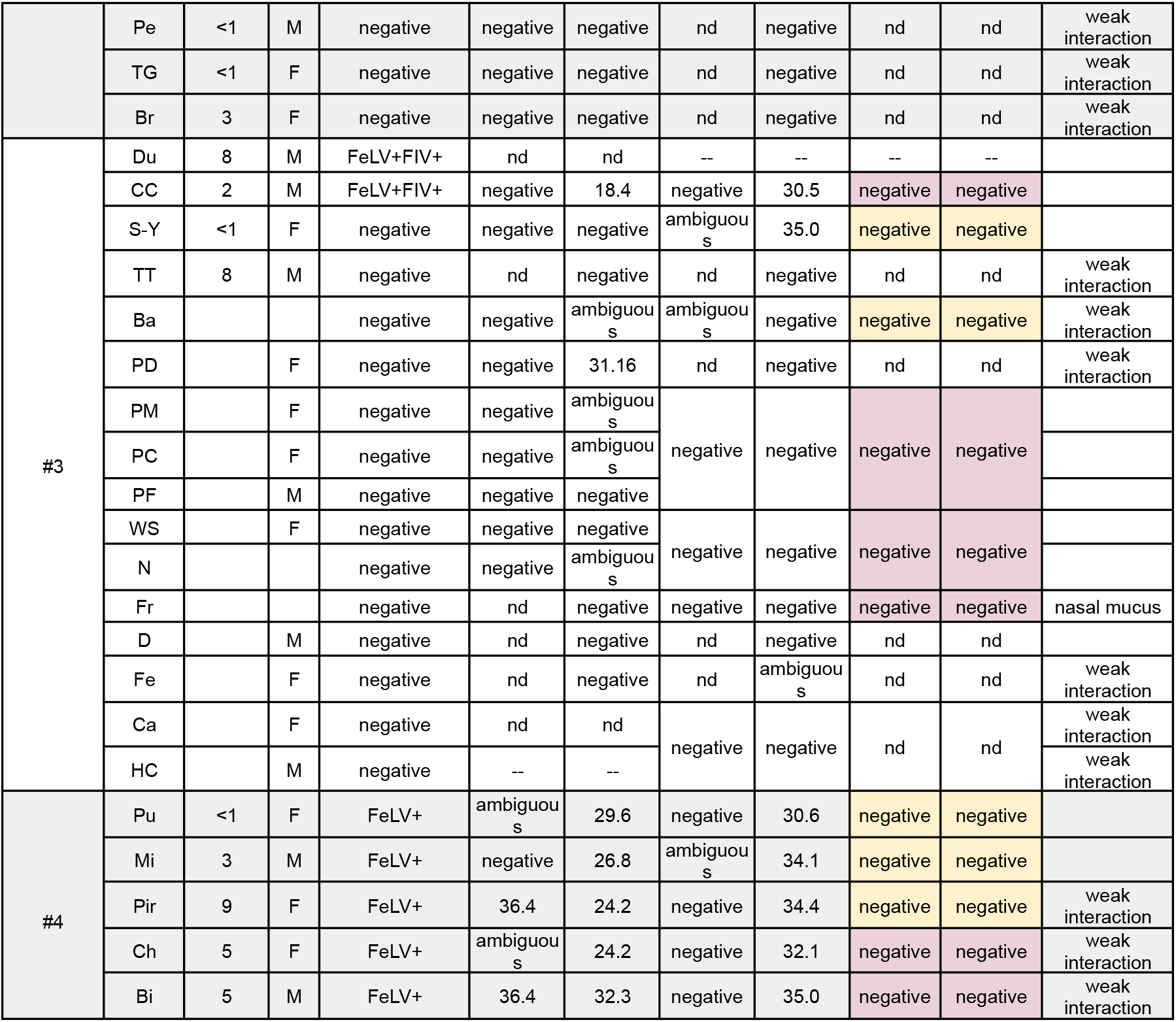
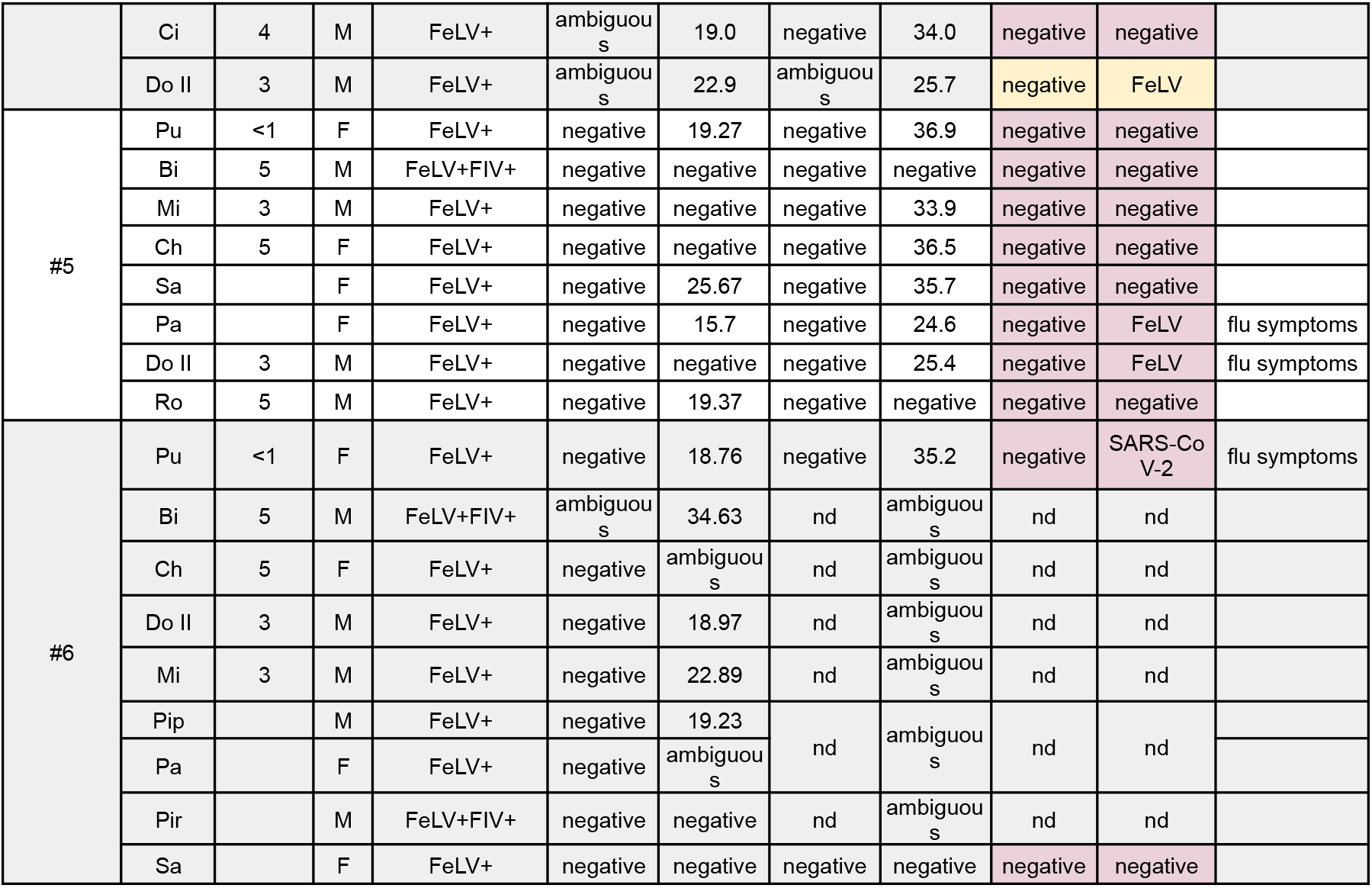
Individual Cq values from cat samples. All cats were tested for presence of FeLV virus and FIV antibodies by analysis of blood (jugular venipuncture) by SNAP test (Idexx™) at admittance in the shelter (“serological status”). RTqPCR of saliva (oral swab or spontaneous licking samples) for FeLV and/or FIV or Nanopore™ metabarcoding (of spontaneous licking samples) were performed and data is listed in the table. Numerical value = mean Cq of RTqPCR duplicates. nd = not done. ambiguous = inconclusive result (one replicate did not amplify OR duplicates amplified only one product of wrong Tm). Yellow highlight = sequencing run 1. Pink highlight = sequencing run 2.

**Table S5.** Results of linear mixed effects model with stepwise elimination examining predictors of variation in Zika virus Cq values (outlier included). Terms included in the best fitting model are listed with significant terms in bold.

| Virus | (Intercept ) | paper | temp | day | paper: temp | paper: day | temp: day | paper: temp:day |
| --- | --- | --- | --- | --- | --- | --- | --- | --- |
| Zika | p = 1.88e-27 | F = 0.11<br>p = 0.741 | F = 0.36<br>p = 0.699 | F = 0.66<br>p = 0.418 | F = 0.44<br>p = 0.647 | **F = 5.45**<br>**p = 0.0219** | **F = 7**<br>**p = 0.00155** | **F = 6.57**<br>**p = 0.00224** |

**Table S6.** Post hoc tests for Zika virus linear mixed effects model with outlier included. Significant terms are listed in bold.

| temp | day | paper | day.trend | SE | df | lower.CL | upper.CL | t.ratio | p.value |
| --- | --- | --- | --- | --- | --- | --- | --- | --- | --- |
| HHH | 10.68966 | W3 | 0.004614 | 0.009551 | 86.04218 | -0.01437 | 0.0236 | 0.48308 | 0.630266 |
| LHH | 10.68966 | W3 | 0.003378 | 0.009551 | 86.04218 | -0.01561 | 0.022365 | 0.35373 | 0.724407 |
| **RT** | **10.68966** | **W3** | **-0.05414** | **0.00966** | **86.04218** | **-0.07335** | **-0.03494** | **-5.60505** | **2.46E-07** |
| HHH | 10.68966 | W54 | 0.010461 | 0.009551 | 86.04218 | -0.00853 | 0.029448 | 1.095292 | 0.276445 |
| LHH | 10.68966 | W54 | -0.00384 | 0.009551 | 86.04218 | -0.02282 | 0.015149 | -0.40175 | 0.688866 |
| RT | 10.68966 | W54 | 0.003358 | 0.009551 | 86.04218 | -0.01563 | 0.022345 | 0.351595 | 0.726002 |

